# A phylogenetic method identifies candidate drivers of the evolution of the SARS-CoV-2 mutation spectrum

**DOI:** 10.1101/2025.01.17.633662

**Authors:** Russ Corbett-Detig

**Affiliations:** Department of Biomolecular Engineering, University of California, Santa Cruz; Genomics Institute, University of California, Santa Cruz

## Abstract

The molecular processes that generate new mutations evolve, but the causal mechanisms are largely unknown. In particular, the relative rates of mutation types (*e*.*g*., C>T), the mutation spectrum, sometimes vary among closely related species and populations. I present an algorithm for subdividing a phylogeny into distinct mutation spectra. By applying this approach to a SARS-CoV-2 phylogeny comprising approximately eight million genome sequences, I identify 10 shifts in the mutation spectrum. I find strong enrichment consistent with candidate causal amino-acid substitutions in the SARS-CoV-2 polymerase, and strikingly three appearances of the same homoplasious substitution are each associated with decreased C>T relative mutation rates. With rapidly growing genomic datasets, this approach and future extensions promises new insights into the mechanisms of evolution of mutational processes.

## Introduction

The molecular machinery that affects the rates and types of new mutations is subject to evolution. In particular, organisms ranging from humans (Harris 2015; Harris and Pritchard 2017) to bacteria (Wei et al. 2022) demonstrate changes in the relative rates of the types of new mutations (*i*.*e*., the mutation spectrum). In mammals the mutation spectrum displays phylogenetic signal such that more closely related species tend to exhibit similar relative mutation rates (Beichman et al. 2023). Associations between changes in the mutation spectrum and specific genetic elements, such as substitutions that affect the function of genome replication enzymes, have the potential to reveal the factors determining the molecular and evolutionary basis of genome maintenance and repair (*e.g., (Robinson et al. 2021; Sasani et al. 2022; Sasani et al. 2024)*). Nonetheless, identifying such associations in natural populations is challenging and there are few cases wherein naturally-occurring causal alleles have been identified.

Recent studies leveraged the vast sequencing dataset for SARS-CoV-2 to investigate differences in the mutation spectrum with unprecedented resolution. Ruis et al. (Ruis et al. 2023) reported a decrease in the rate of G>T mutations in Omicron and descendant lineages. Similarly (Bloom et al. 2023) corroborated this finding and discovered additional shifts in the mutation spectrum across SARS-CoV-2 lineages since its emergence in human populations. However, these and related studies generally rely on comparisons of mutation spectra using *a priori* subdivisions of the samples (e.g., clades or populations). This means that if the mutation spectrum changes are not associated with those specific subdivisions they may be overlooked and the causal mechanisms difficult to discern.

Several phylogenetic methods identify changes in the mutation process across lineages (*e.g., (Huelsenbeck et al. 2000; Jayaswal et al. 2014)*). For example, (Blanquart and Lartillot 2006) developed a Bayesian method that can identify branches along the phylogeny where the substitution process changed. In principle, these approaches could facilitate identification of the molecular drivers of mutation spectrum evolution. However, existing maximum likelihood and Bayesian frameworks are limited due to computational demands, and new approaches that can effectively leverage vast genomic datasets are essential for comprehensively exploring the evolution of molecular evolutionary processes.

## Results and Discussion

### Algorithm Overview

I introduce spectrumSplits, a method for subdividing a phylogeny into non-overlapping subtrees with distinct mutational spectra. This approach accepts a phylogenetic tree where mutations have been assigned to internal nodes via maximum parsimony using UShER (Turakhia et al. 2021). The algorithm performs a depth-first pre-order traversal of the phylogeny and compares the mutation spectra for the subtrees on either side of a given node using a X^2^ test statistic where the categories are the counts of each mononucleotide mutation type (*i.e*., A>C, A>G, etc.) compared between the two subtrees. If the maximum recorded X^2^ is greater than a specified threshold, the tree is bisected at the corresponding node, resulting in a “spectrum split”, and the procedure is repeated on the two resulting subtress until no additional nodes are discovered that exceed the threshold. The first split is therefore performed at the node with the largest X^2^ identified in the entire phylogeny. This is similar to the automated lineage designation method autolin (McBroome et al. 2024). I note that this procedure assumes that the evolution of the mutation spectrum is discontinuous and may not be ideal for gradual changes.

### Pseudocode

initialize treeVector with tree

while length(treeVector) > 0:

for t in treeVector:

for node in t:

compute X^2^

if max X^2^ > threshold

bisect t at max X^2^ node into t1, t2

add t1, t2 to treeVector

delete t from treeVector

### Spectrum partitions

I applied spectrumSplits to the comprehensive public SARS-CoV-2 phylogeny from the UShER project (dated 9/21/2024, (McBroome et al. 2021; Turakhia et al. 2021)). After obtaining the phylogeny, which receives substantial curation but has different goals than the present analysis (Hinrichs et al. 2023), I applied conservative methods to remove potentially spurious samples and mutations (see Methods). The tree used in analysis contained 7,593,765 samples with 4,668,537 mutations and is available for [interactive visualization via taxonium](https://taxonium.org/?protoUrl=https%3A%2F%2Fhgwdev.gi.ucsc.edu%2F%7Eangie%2Fpublic-2024-09-28.all.masked.prunedJunk.prunedSites.jsonl.gz) (Sanderson 2022; Kramer et al. 2023). I identified 10 spectrum splits associated with changes in the mutation spectrum. This procedure ran with 1000 bootstraps on 100 threads in 68 minutes.

### Robustness of spectrumSplits

I used nonparametric bootstraps where I resampled alignment columns with replacement to evaluate the robustness of spectrumSplits using three metrics. First, I computed the proportion of times that a spectrum split in the full dataset is recovered in each bootstrap. Second, because a spectrum split node might be similar but not identical to a bootstrap node, I calculated the mean distance, in number of edges, between each spectrum split identified in the full dataset and the nearest bootstrap spectrum split. Third, I calculated the maximum Jaccard set similarity of the node set descendant from a spectrum split between full and bootstrapped outputs. For each spectrum split, I found that bootstrap support is moderate to high (median 0.61, range 0.3–1.0, Table 1), that the distance to the nearest bootstrap split is low (median 0.72 edges, range 0.0–3.01), and furthermore that the Jaccard similarity is high (median 0.96, range 0.68–1.0). I conclude that the selection of spectrum splits is reasonably robust.

**Table 1.**
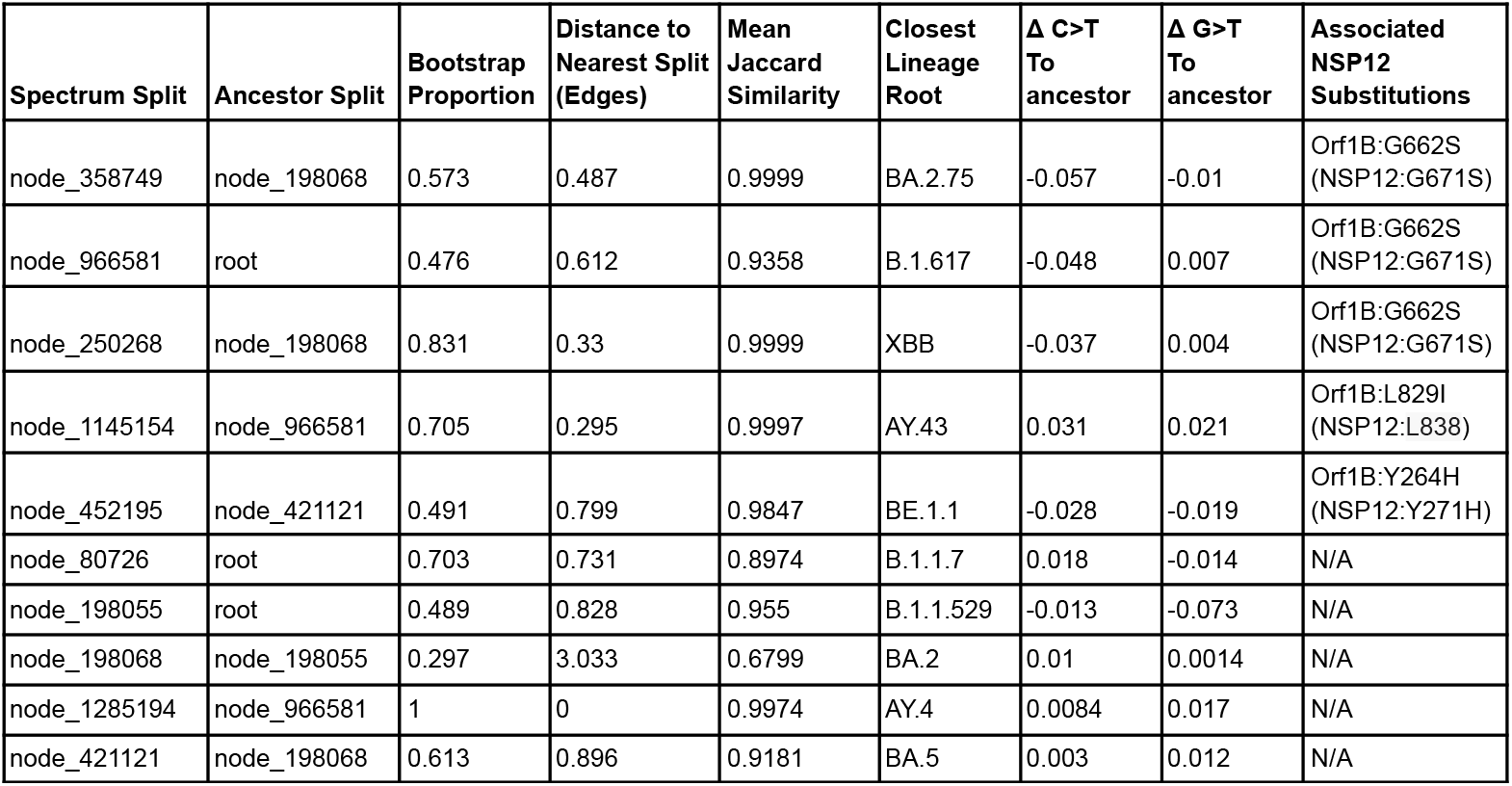
Summary of each spectrum split. This includes the ancestor spectrum split (or root, as applicable), the proportion of bootstraps that recovered the same node, the distance to the nearest spectrum split identified in bootstraps, the mean maximum Jaccard set similarity between the bootstrapped spectra and split in the full dataset, the closest PANGO lineage root (O’Toole et al. 2021) to each spectrum split, the changes in the C>T and G>T relative mutation rates compared to the ancestor of that spectrum split and the amino acid substitutions in the polymerase in Orf1B coordinates and NSP12 coordinates in parentheses. The table is sorted by |Δ C>T|.

### Primary Components of Mutation Spectrum Evolution

A principal component analysis identified the major axes of variation as the relative rates of C>T and G>T mutation. Just two principal components (PCs) explain most of the variation among mutation spectra (78% and 21%, Figure 1A-C). The first is largely related to the relative rate of G>T mutation, and stratification along this PC subdivides omicron and non-omicron lineages (Figure 1B-D), implying that a large shift in the relative rate of G>T evolution evolved once in SARS-CoV-2. The second is driven by variation in the C>T mutation rate which is more variable across the phylogeny (Figure 1B,C,E). My results are consistent with (Bloom et al. 2023; Ruis et al. 2023), and may also reveal causal mechanisms not previously described.

**Figure 1.**
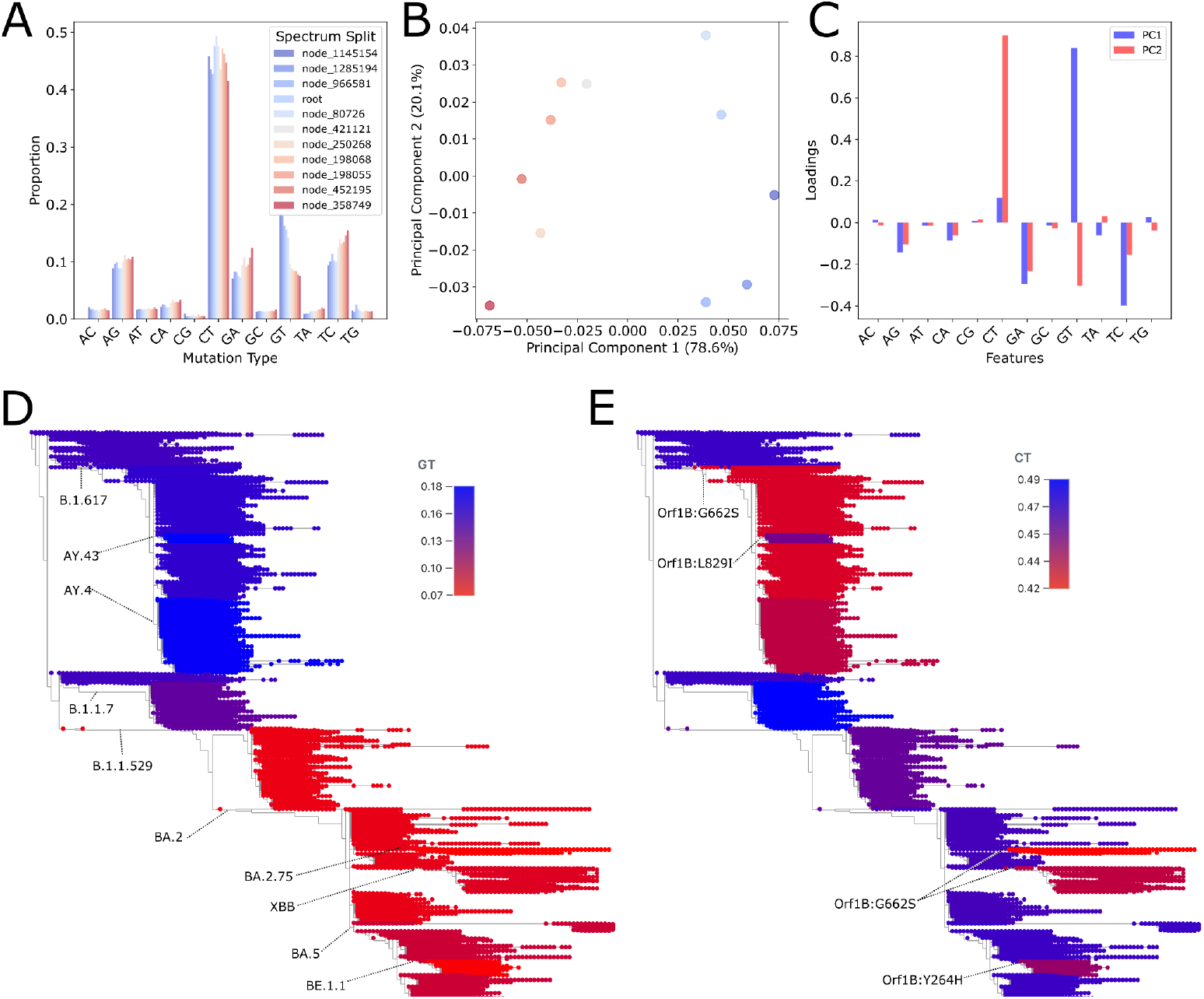
Results from Spectrum Splits. (A) Relative proportion of each mutation type in each identified spectrum including the root. (B) Principal component analysis of the 11 mutation spectra obtained from this analysis, colored as in panel A. (C) PCA loadings for the first two principal components across the 12 relative mutation rates. (D) Estimated relative rates of G>T substitution in each spectrum split. The nearest PANGO lineage root for each spectrum split is labelled. (E) Estimated relative rates of C>T substitution in each spectrum split. Amino acid substitutions in the viral polymerase are labeled at the nodes where they occurred. An [interactive visualization is available here](https://taxonium.org/?protoUrl=https%3A%2F%2Fhgwdev.gi.ucsc.edu%2F%7Eangie%2Fpublic-2024-09-28.all.masked.prunedJunk.prunedSites.jsonl.gz).

### Candidate Causal Mutations of Spectrum Splits

I identified a set of candidate causal mutations by extracting amino acid substitutions in the viral polymerase protein, NSP12, that occur near to spectrum splits. In light of the bootstrapping results, I included mutations at nodes wherein the descendant set has a substantial overlap with the spectrum split (see Methods). Among ten spectrum splits, five are associated with a non-synonymous substitution in NSP12 (Table 1; P < 0.0001, Permutation Test). Furthermore, the five spectrum splits associated with non-synonymous substitutions in the polymerase, are those with the largest change in the C>T relative mutation rate (0.028–0.057 vs. 0.003–0.018; P = 0.0079, Mann-Whitney U Test). This suggests that substitutions within the polymerase contribute to mutation spectrum evolution in SARS-CoV-2.

The association between decreased C>T mutation rates and recurrent substitutions of Orf1B:G662S (NSP12:G671S) suggests it may be a causal mutation of mutation spectrum evolution. Orf1B:G662S enhances replication possibly by contributing to the stability of the replication complex (Kim et al. 2023) and may enhance viral fitness. I found this substitution in association with the three largest decreases in the relative rate of C>T mutation among all spectrum splits (−0.037, -0.048, and -0.057; P = 0.0167, Mann Whitney U Test). In fact, these are the only nodes where Orf1B:G662S evolved with more than 10,000 descendant samples, and it is unlikely that three randomly-selected nodes would be associated with the same amino acid substitution (P < 0.0001, Permutation Test). Therefore, this mutation is a strong target for future functional investigations of mutation spectrum evolution. Furthermore, I speculate that C>T mutation rate evolution might provide a proxy for polymerase function (either fidelity or speed) when comparing viral lineages and therefore present a valuable metric for functional surveillance of SARS-CoV-2 to identify epidemiologically distinct lineages.

That there are no amino-acid altering substitutions in NSP12 associated with the origins of Omicron (B.1.1.529), which displays one of the largest changes in mutation spectrum, suggests that distinct factors might drive the evolution of C>T and G>T relative mutation rates. In particular, mutations in other parts of the genome could affect the mutation spectrum. For example, ORF1B:I1566V (NSP14:I42V) in the viral proofreading domain is a strong plausible candidate (Bloom et al. 2023). Nonetheless, Omicron also shifted towards a replication niche in the upper lungs (Hui et al. 2022) which might affect exposure to mutagens and impact the mutation spectrum (Ruis et al. 2023). In the absence of replicate large changes in the relative G>T mutation rates among SARS-CoV-2 lineages, distinguishing among these possibilities is difficult. Future sampling and application of spectrumSplits could help identify which additional factors are responsible for mutation spectrum evolution.

## Conclusion

SpectrumSplits rapidly identifies candidate nodes that demonstrate changes in their mutation spectrum, which can then be associated with candidate mutations for downstream experiments and has the potential to empower surveillance of pathogen functional evolution. In particular, the discovery of a strong association between Orf1B:G662S and decreased C>T mutation rates is an appealing future direction for characterizing the functional basis of the evolution of the mutation spectrum. As genomic datasets are rapidly growing in size, spectrumSplits will also enable investigations of the factors that affect evolutionary processes across diverse organisms. Although this method is currently designed for a non-recombining phylogeny, extensions using ancestral recombination graphs (Ethier and Griffiths 1990; Wong et al. 2024) that consider both the local non-recombined gene tree surrounding a spectrum-affecting mutation and unlinked gene trees is possible. Similarly, it is possible and it could be informative to extend this approach to identify evolution of mutation processes using additional features such as trinucleotide context and multi-nucleotide mutations. Therefore, spectrumSplits will provide a powerful basis for investigating the causes of the evolution of the mutation processes.

## Methods

### Quality Control

Sequencing and assembly errors have been widely documented in SARS-CoV-2 (*e.g., (Turakhia et al. 2020; De Maio et al.)*) and if unaddressed might lead to spurious inferences of changes in the mutation spectrum. I used two approaches to mitigate their impacts. First, I pruned subtrees where the ratio of mutations:tips exceeded 3, an effect commonly observed with amplicon-dropout and resulting assembly errors. Second, I masked individual positions whose relative mutation rates increased dramatically at a particular node in the phylogeny — using an approach that is similar to spectrumSplits but considers mutations at one position vs all other mutations. I visually inspected the masking results to determine if mutations excluded were likely to be spurious. The set of masked nodes and mutations are available in the github repository associated with this project.

### Spectrum Splits

I applied spectrum splits to identify nodes where the descendant mutation spectrum changed relative to its ancestor. I consider all mutation classes because SARS-CoV-2 is a plus-stranded RNA virus and the strand is an important determinant of the mutation spectra (e.g., (Simmonds 2020; Turakhia et al. 2020)), rather than collapse mutation types by reverse complementarity. I ran spectrumSplits requiring a minimum X^2^ test statistic of 500. Determining the exact number of statistical tests implied is not straightforward — *i.e*., the algorithm traverses the tree or resulting subtrees repeatedly, many nodes are evaluated in each traversal, and the number of traversals is not known in advance. Nonetheless, the X^2^ value used is conservative as the final procedure required fewer than 5 million statistical tests.

### Bootstrapping

I used non-parametric bootstrapping to resample alignment columns to quantify the robustness of inferred spectrum splits. I compared bootstrapped datasets with the full dataset in terms of spectrum split identification, phylogenetic proximity to other spectrum splits, and the Jaccard set similarity in the descendant nodes contained within a given spectrum split and its maximally similar bootstrap split.

### Permutation Tests

To determine if there is an enrichment for non-synonymous mutations in the NSP12 associated with spectrum splits, I annotated each node with mutations found at the same node or at adjacent nodes that shared more than 99.99% of the descendant samples. For instance, node_966581 (B.1.617, Delta) includes a few additional samples that apparently contain subsets of the Delta-specific substitutions. I then performed permutation by randomly resampling nodes from the full phylogeny with a minimum of 10,000 descendants and identifying associated non-synonymous polymorphisms in NSP12, and then determining if this permuted set of nodes is associated with the same or more NSP12 amino-acid altering substitutions than the true spectrum splits. I performed an equivalent test for association of spectrum splits with a single non-synonymous substitution.

## Software and Data Availability

SpectrumSplits and ancillary methods are available from https://github.com/russcd/spectrumSplits. The phylogeny used for this analysis is available at http://hgdownload.soe.ucsc.edu/goldenPath/wuhCor1/UShER_SARS-CoV-2/2024/09/28/public-2024-09-28.all.masked.pb.gz.

## Acknowledgements

I thank Alex Ioannides, Robert Lanfear, Shelbi Russell, Yatish Turkahia, Angie Hinrichs, Landen Gozashti, Pratik Katte, Erik Enbody and members of the Corbett-Detig and Turkahia labs for helpful discussions. I gratefully acknowledge all individuals and groups who sequenced and shared SARS-CoV-2 genome sequences. Funding was provided by NIH/NIGMS R35GM128932.

